# Concurrent Stereotactic Body Radiation Therapy and KRAS Inhibition Synergistically Improve Pre-clinical Pancreatic Cancer Treatment

**DOI:** 10.64898/2026.07.10.737883

**Authors:** Tianyu Wang, Liang Wang, Jiaqian Xu, Youming Guo, Ling Xia, Yuting Li, Fada Guan, Boyi Gan, David S. Hong, Vincent Bernard, Dadi Jiang, Albert C. Koong

## Abstract

Pancreatic ductal adenocarcinoma (PDAC) is one of the most challenging cancers to treat due to the dismal survival rate, poor post-treatment outcome and profound resistance to a wide range of therapies. With mutant *KRAS* being a key driver, small molecule inhibitors targeting KRAS or pan-RAS (KRASi) have demonstrated exciting preclinical and early clinical anti-tumor efficacy, and the pan-RAS(ON) inhibitor daraxonrasib (RMC-6236) recently achieved Phase 3 clinically meaningful improvements in patient survival compared to chemotherapy. But resistance to RAS/KRAS inhibitor inevitably develops, which limits and compromises the treatment outcome. In this study, we investigated the combination of stereotactic body radiation therapy (SBRT) and KRAS inhibition (MRTX1133 and daraxonrasib) in the treatment of preclinical PDAC models. We found that this combination strategy synergistically suppresses PDAC cell growth *in vitro* and enhances tumor control while minimizing local recurrence in orthotopically implanted KPC (*LSL-Kras*^*G12D/+*^*;Trp53*^*R172H/+*^*;Pdx1-Cre*) murine PDAC tumors *in vivo*. As radiation therapy (RT) induces ferroptosis in multiple cancer types and mutant KRAS promotes various anti-ferroptotic mechanisms, we tested the role of ferroptosis in promoting tumor-control efficacy. Intriguingly, the addition of a ferroptosis inhibitor, liproxstatin-1, to the combination therapy significantly abrogated the *in vivo* synergism between SBRT and KRAS inhibition, suggesting that treatment-induced ferroptosis at least partially drives the synergistic efficacy of this combination strategy. Our study indicates that this SBRT-KRASi combination has the potential to overcome treatment resistance and improve outcomes in PDAC patients. These data directly support the design of a planned multi-center Phase 2 clinical trial with this combination strategy in locally advanced PDAC.

## INTRODUCTION

Pancreatic ductal adenocarcinoma (PDAC) is the 3^rd^ most common cause of cancer-related death in the U.S. and predicted to be the 2^nd^ cause of cancer death by 2030^1,2^. In contrast to the steadily improving survival rate for most cancers, the 5-year relative survival for patients with PDAC remains at 13% and has only modestly increased in the last few decades^1,3,4^. The majority of PDAC cases are diagnosed at an advanced stage, making curative treatment difficult, thus leading to a poor overall prognosis^5^. Therefore, the development of targeted therapies based on a greater understanding of the molecular vulnerabilities of PDAC is urgently needed.

Despite significant improvements in the basic understanding of PDAC, surgical resection remains the primary option for extended survival, but only a small fraction (10-15%) of patients present with resectable disease. Patients with locally advanced pancreatic cancer (LAPC) are not amenable to resection and account for 30-40% of all cases of PDAC. Chemotherapy is modestly effective, but often fails to control the primary tumor^6^ and local progression often directly leads to morbidity and mortality^7^. For non-resectable PDAC, radiation therapy (RT) contributes primarily to local control. Definitive chemoradiation to PDAC is challenging because the adjacent stomach and duodenum cannot tolerate high doses of RT^8^. In 2004, we were the first to report a prospective dose escalation study in pancreatic cancer using a highly conformal technique called stereotactic body radiation therapy (SBRT) or stereotactic ablative radiotherapy (SABR) that utilizes advanced image guidance to deliver high doses of radiation to the tumor^9^. Today, SBRT is an acceptable first-line treatment option for certain patients with LAPC according to NCCN (v2.2024) and ASTRO guidelines^10^.

Genomic analyses reveals that the majority (70-95%) of PDAC cases have an activating point mutation of the *KRAS* proto-oncogene at codon 12 (glycine, G)^11-14^. These single-nucleotide mutations lead to changes including G12D (40%) then followed by G12V (33%). The *KRAS* gene encodes a small GTPase that serves as a molecular switch for various cellular signaling processes by coupling cell surface growth factor receptor with transcription factor activation^12,13^. Once activated via binding to GTP, KRAS activates multiple downstream signaling pathways, including RAF-MEK-ERK and PI3K-AKT-mTOR, further activating transcription factors like ELK, JUN and MYC to stimulate cellular differentiation, proliferation, survival, adhesion, migration and transformation^15^. An activating point mutation of KRAS abrogates its intrinsic GTPase activity and locks KRAS at GTP-bound (active) status, thus constitutively activating downstream signaling pathways.

Ferroptosis is an iron-dependent form of regulated cell death that is induced by excessive lipid peroxidation^16-18^. Polyunsaturated fatty acids (PUFAs) in the phospholipid component of cellular membranes are susceptible to lipid peroxidation in iron- and oxygen-rich cellular environments, which, if left unchecked, can damage membrane integrity and induce ferroptotic cell death^19^. Cells have evolved elegant defense mechanisms against ferroptosis to eliminate harmful lipid peroxides, including the cystine uptake-coupled SLC7A11-glutathione-GPX4 signaling axis as a major component^20-22^ and GPX4-independent mechanisms such as the conversion of ubiquinone (CoQ) to ubiquinol (CoQH_2_), a lipophilic radical-trapping antioxidant, by ferroptosis suppressor protein 1 (FSP1) and dihydroorotate dehydrogenase (DHODH)^23-25^. Ferroptosis is recognized as a natural tumor suppression mechanism, and is highly relevant to cancer therapeutic strategies^26^. Recent studies showed that RT can potently induce ferroptosis suggesting that pharmacological ferroptosis inducers (FINs) may be combined with RT to improve radiosensitivity and overcome radioresistance^27-30^.

Targeting ferroptosis is a promising therapeutic strategy for PDAC treatment based on recent studies^31^. Pancreas-specific deletion of *Slc7a11* in the *Kras*^*G12D/+*^*;Trp53*^*R172H/+*^; *Pdx1-Cre*^*+*^ (KPC) mice induced tumor-specific ferroptosis and abolished pancreatic tumor development through limiting CoA and GSH synthesis downstream of cystine uptake. The same effect can be replicated with administration of cyst(e)inase that depletes cysteine and cystine *in vivo*^32^. Inhibition of glutamic-oxaloacetic transaminase 1 (GOT1), an enzyme involved in maintaining redox balance, represses mitochondrial metabolism and increases labile iron availability through autophagy, thereby enhancing PDAC sensitivity to ferroptosis induction^33^.

Although KRAS was previously thought to be an undruggable target^34^, the recent success of KRAS^G12C^ inhibitors sparked considerable enthusiasm in developing other inhibitors that directly target KRAS^35,36^. Various allele-specific as well as pan-KRAS/pan-RAS inhibitors/degraders are at various stages of development^37^. MRTX1133^38,39^ is a noncovalent inhibitor of KRAS^G12D^ with prominent anti-cancer activities in preclinical models of KRAS^G12D^-driven lung, pancreatic and colorectal adenocarcinoma^40-43^. Daraxonrasib (RMC-6236), a first-in-class RAS(ON) inhibitor designed to target multiple active RAS isoforms, has demonstrated highly promising clinical progress across both early- and late-stage trials. In the recent Phase 3 RASolute 302 trial evaluating 2^nd^ line treatment of patients with metastatic PDAC, daraxonrasib achieved statistically significant and clinically meaningful improvements in progression-free survival (PFS) and overall survival (OS) compared with standard-of-care chemotherapy^44^. Although initially efficacious, tumors inevitably develop resistance to KRAS inhibitors as single treatment modality and strategies to overcome treatment resistance are critical for maintaining clinical efficacy^45^. Both intrinsic and acquired resistance to KRAS inhibitors are driven by genetic and non-genetic mechanisms to reactivate upstream RTKs, KRAS itself, or their effector pathways^46,47^. These findings suggest that KRAS/pan-RAS inhibitors alone will not be sufficient to overcome adaptive mechanisms leading to resistance, therefore combinational strategies will be needed to improve overall treatment outcome.

## MATERIALS AND METHODS

### Cell culture

The mouse KPC0 cell line is a gift from Dr. Anirban Maitra (NYU) and originally derived from the *LSL-Kras*^*G12D/+*^*;Trp53*^*R172H/+*^*;Pdx1-Cre*^*+*^ tumor model. The fast-growing variant, FG, of human pancreatic cell line COLO 357 was previously described^48^. Capan-2 and MIA Paca-2 cells were purchased from the American Type Culture Collection. Panc-1 cells were purchased from Sigma Aldrich. The human PDAC PDX-derived PATC153 cell line was distributed by the Translational Research to AdvanCe Therapeutics and Innovation in ONcology (TRACTION) platform at the UT MD Anderson Cancer Center^49^. All cell lines were free of mycoplasma at the time of the assay tested with the MycoAlert Mycoplasma Detection Kit (Lonza). No cell line used in this study has been found in the International Cell Line Authentication Committee (ICLAC) database of commonly misidentified cell lines (version 11). Cell lines were cultured in Dulbecco’s modified Eagle’s medium (DMEM) with 10% (volume/volume; v/v) heat-inactivated FBS (Gibco) and 1% (v/v) penicillin/streptomycin (Gibco) at 37 °C and 5% CO_2_.

### Stable cell line generation through viral transduction

KPC0 cells overexpressing firefly luciferase (KPC0-luc) were generated as reported previously^50^. Briefly, HEK293T packaging cells were transfected with pLEX-luciferase lentiviral vector (Thermo Scientific) together with pCMV-VSVG and pCMV-Δ8.2 lentiviral packaging plasmids. 24 and 48 h later, viral supernatant was collected and filtered through a 0.45 μm filter. To infect the target cell lines, 0.8 μg/ml polybrene (Sigma Aldrich) was added to the viral supernatant before adding to the target cells. 24 h after starting infection, 1-5 μg/ml puromycin (Gibco) was added to the cells for antibiotic selection. Luciferase expression was verified with Bright-Glo Luciferase Assay (Promega).

### *In vitro* radiation – inhibitor synergism assay

Cells were seeded in 96-well plates at a density of 1,000 – 2,000 cells per well. After overnight incubation at 37ºC in a CO_2_ incubator, the cells were pre-treated with DMSO (vehicle), MRTX1133, or RMC-6236 at indicated concentrations for 4 hrs prior to irradiation. Radiation was delivered using a cabinet X-ray irradiator (X-RAD 320, Precision X-Ray) at a dose rate of approximately 250 cGy/minute. When the vehicle-treated wells reached 100% confluency (approximately 96 hrs post-treatment), cell density was determined using crystal violet staining and quantitated using a plate reader (BioTek Epoch). Synergy between radiation and KRAS inhibitor was determined using the SynergyFinderPlus R package^51^ with the Highest Single Agent (HSA) model^52^.

### Lipid peroxidation assay

To measure lipid peroxidation, *KRAS/Kras*-mutant human or mouse PDAC cells were seeded in 6-well plates at a density of 5 x 10^5^ cells/well and assigned to the following treatment groups: vehicle (DMSO) control, MRTX1133 (200 nM) or RMC-6236 alone (100 nM), RT alone (10 Gy), MRTX1133 or RMC-6236 + RT, and MRTX1133- or RMC-6236 + RT + liproxstatin-1 (10 μM]) after overnight incubation. The treatment with compound(s) lasted for 24 hrs and irradiation was performed 6 hrs before the end of the compound treatment.

After treatment, cells were stained with 2 μM BODIPY 581/591 C11 (Thermo Fisher Scientific) dissolved in DPBS for 15 min at 37°C in the dark. The cells were then harvested using TrypLE cell dissociation reagent, washed with DPBS, and resuspended in DPBS. Flow cytometry was performed using an Attune NxT Flow Cytometer (Thermo Fisher Scientific) according to the manufacture recommended operating procedures. Lipid peroxidation was quantified by measuring fluorescence intensity from the oxidized state (BL1 channel).

### Orthotopic tumor cell injection into mouse pancreas

All animal procedures were approved by the Institutional Animal Care and Use Committee (IACUC) of the UT MD Anderson Cancer Center. KPC0-luc cells stably expressing luciferase cultured in 10-cm tissue culture plates were harvested using TrypLE cell dissociation reagent (Gibco), washed and resuspended in Dulbecco’s Phosphate-Buffered Saline (DPBS) at a density of 2.5 x 10^5^ cells per 20 μL.

8∼12-week-old C57BL/6 mice were anesthetized using isoflurane (2 - 3% induction, 1.5 - 2% maintenance) and placed in a lateral recumbent position. Pre-operative analgesia was administered 30 min prior to the surgical procedure via a subcutaneous injection of buprenorphine extended-release (Buprenorphine-ER, 1 mg/kg). A small surgical incision was made in the left subcostal abdomen to expose the spleen and the tail of the pancreas. The tumor cell suspension was slowly injected into the pancreatic tail using a 28-gauge insulin syringe (BD Biosciences).

Immediately following needle withdrawal, a titanium ligating microclip (Weck Horizon) was placed 1 mm inferior to the injection site to serve as a fiducial marker for subsequent microCT-guided treatment planning on the Small Animal Radiation Research Platform (SARRP, Xstrahl). The pancreas was gently returned to the abdominal cavity. The abdominal wall and peritoneum were closed using 4-0 absorbable sutures (Ethicon), and the skin incision was closed with surgical wound clips. A second maintenance dose of Buprenorphine-ER was administered 48 hrs post-surgery to ensure sustained post-operative pain management. On Day 10 post-surgery, tumor engraftment was confirmed via bioluminescence imaging using an In Vivo Imaging System (IVIS, Caliper Life Sciences) in the Small Animal Imaging Facility (SAIF) at the UT MD Anderson Cancer Center. Upon confirmation of tumor establishment, the tumor-bearing mice were subsequently randomized into treatment groups.

### Micro-CT guided small animal SBRT

On Day 11 post-surgery, mice assigned to the SBRT-alone or SBRT + KRASi combination treatment group were anesthetized via isoflurane inhalation and secured on the robotic positioning couch of the SARRP. The orthotopic tumor was localized using SARRP’s integrated micro-computed tomography (micro-CT) imaging system based on the location of the titanium ligating microclip on the CT image for precise target alignment.

The radiation treatment isocenter was mapped directly to the tumor coordinates. Targeted X-ray irradiation was delivered using an SBRT-like treatment planning featuring a 4-beam configuration (4 angles) at a dose rate of approximately 250 cGy/min. 8 Gy was administered every 24 hrs for 5 consecutive days with a total cumulative dose of 40 Gy per mouse. Following each radiation fraction, the treated mice were placed in a temperature-controlled recovery cage over a heating pad until they fully regained consciousness and mobility before returning to the original housing cage.

### KRAS inhibitor (KRASi) administration

The KRASi MRTX1133 and RMC-6236 were provided by TRACTION or purchased from a commercial vendor (Chemietek) with purity confirmed by HPLC. For groups receiving KRASi alone or in combination with SBRT, the compounds were administered as follows: MRTX1133 was delivered intraperitoneally (*i*.*p*.) twice daily (*b*.*i*.*d*.) at a dose of 30 mg/kg, and RMC-6236 was administered via oral gavage (*p*.*o*.) once daily (*q*.*d*.) at a dose of 25 mg/kg. For the SBRT + KRASi + liproxstatin-1 group, liproxstatin-1 (Cayman Chemical) was administrated at a dose of 10 mg/kg together with KRAS inhibitor daily for the first 7 days then once every 2 days afterwards. Vehicles used for dissolving the compounds are 50 mM citric acid, pH 5.0 with 10% Captisol (m/v) for MRTX1133, 10/20/10/60% (v/v) DMSO/PEG 400/Solutol HS15/water for RMC-6236, and 18% Captisol (m/v) with 10% DMSO (v/v) in saline for liproxstatin-1.

11 days after tumor cell implantation, the initial dose of KRASi was administered 6 hrs before the first fraction of SBRT. For the KRASi-alone and the SBRT + RMC-6236 groups, KRASi was administered continuously until the mice reached predetermined institutional humane endpoints. As mice in the SBRT + MRTX1133 group demonstrated complete and sustained tumor regression with no observed recurrence, MRTX1133 administration in this group was stopped when the last animal in the MRTX1133-alone group reached its humane endpoint.

## RESULTS

To assess whether the addition of KRAS inhibition may sensitize PDAC cells to RT, we first performed an *in vitro* cell proliferation assay in a 96-well format. In this assay, increasing RT doses (0 – 10 Gy with 2 Gy increment) are applied to the columns along the *x*-axis and increasing KRASi doses are applied to the rows along the *y*-axis, which forms an RT-KRASi treatment matrix that allows the evaluation of synergy between these two types of treatment. As shown in **Fig. 1**, both mouse and human PDAC cells harboring various *Kras/KRAS* mutations, including KPC0 (*Kras*^*G12D*^), Panc-1 (*KRAS*^*G12D*^), FG (*KRAS*^*G12D*^), PATC153 (*KRAS*^*G12V*^), Capan-2 (*KRAS*^*G12V*^) and MIA PaCa-2 (*KRAS*^*G12C*^), displayed synergy between RT and either allele-specific (MRTX-1133, KRAS^G12D^ inhibitor) or pan-RAS(ON) inhibitor (RMC-6236/daraxonrasib). These different PDAC cell lines displayed diverse sensitivity profiles to single treatment, with KPC0 being relatively radioresistant (**Fig. 1A**) while Panc-1 and FG were relatively resistant to KRASi (**Fig. 1B and C**). Accordingly, in KPC0 cells the synergy scores determined by the Highest Single Agent (HSA) model increased mainly along the RT dose axis at relatively low KRASi doses (**Fig. 1A**), while in Panc-1 and FG cells the synergy scores increased mainly along the KRASi dose axis at relatively low RT doses (**Fig. 1B and C**). Interestingly, we found that the *KRAS*^*G12C*^-bearing MIA Paca-2 cells are significantly more sensitive to RMC-6236 compared to AMG-510, a KRAS^G12C^-specific inhibitor and low dose RT sensitized these cells to both RMC-6236 and AMG-510 (**Fig. 1F**). Taken together, this panel of PDAC cell lines, representing varying degrees of intrinsic resistance to RT or KRASi alone across a diverse *KRAS* mutation spectrum, demonstrated that consistent synergy between the two treatments in suppressing PDAC cell growth.

**Figure 1.**
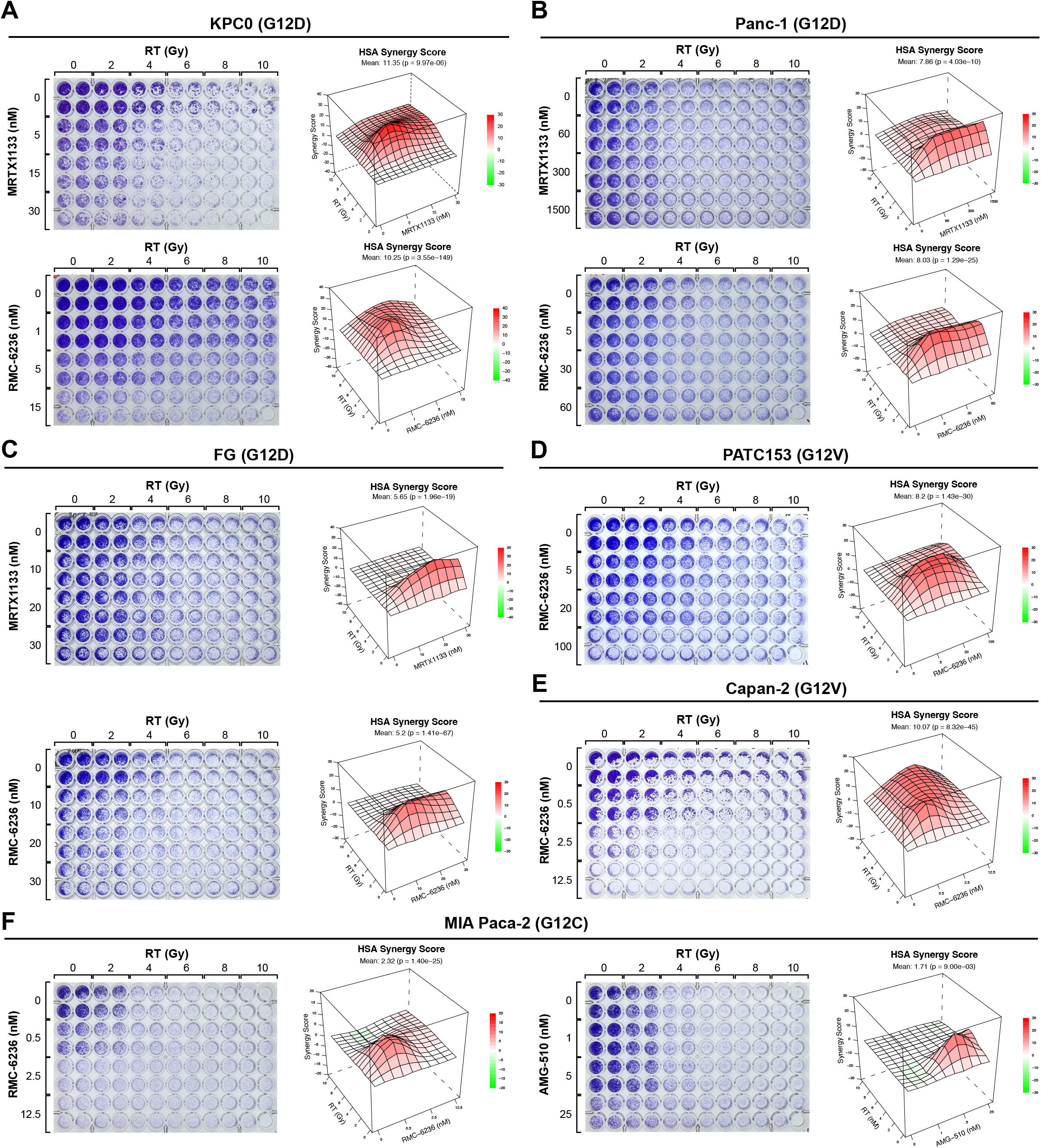
*In vitro* radiation-KRAS inhibitor synergy assay. The synergistic effect of radiation combined with KRAS inhibition was evaluated in a 96-well cell growth format. In this format, radiation doses are distributed along the *x*-axis (0-10 Gy) and KRAS inhibitor doses are along the *y*-axix (variable depending on cell line). Mouse (**A**) and human (**B-F**) PDAC cell lines with different *Kras/KRAS* mutations, including G12D (**A-C**), G12V (**D-E**) and G12C (**F**) as shown in the panel title. The cells were pre-treated with KRAS inhibitors for 4 hours before irradiation and the assay stopped when cells in the vehicle-only control group reached confluency, followed by crystal violet staining and quantification. The image of the stained plate is shown on the left side of each panel. The synergy between radiation and drug was determined with the Highest single agent (HSA) model and the 3D Surface Plot of the synergy model is shown on the right side of the panel.

To determine whether the combination of RT and KRASi further enhances lipid peroxidation as driver of ferroptosis in PDAC cells, we performed BODIPY 581/591 C11 staining and flow cytometry analysis. Consistent with previous findings in other cancer types^27-30^, RT robustly enhanced lipid peroxidation in KPC0 and FG cells. Interestingly, KRASi alone, with either allele-specific (MRTX1133, **Fig. 2B**) or pan-RAS(ON) (RMC-6236, **Fig. 2A and C**) inhibitor, also significantly increased lipid peroxidation to an even greater degree compared to RT (**Fig. 2**), suggesting that oncogenic KRAS signaling actively suppresses lipid peroxidation and dampens PDAC cell sensitivity to ferroptosis. When RT is combined with KRASi, lipid peroxidation was further enhanced (**Fig. 2**). The addition of ferroptosis inhibitor liproxstatin-1 efficiently suppressed the enhanced lipid peroxidation by RT + KRASi treatment, confirming that RT + KRASi enhances ferroptosis induction in PDAC cells.

**Figure 2.**
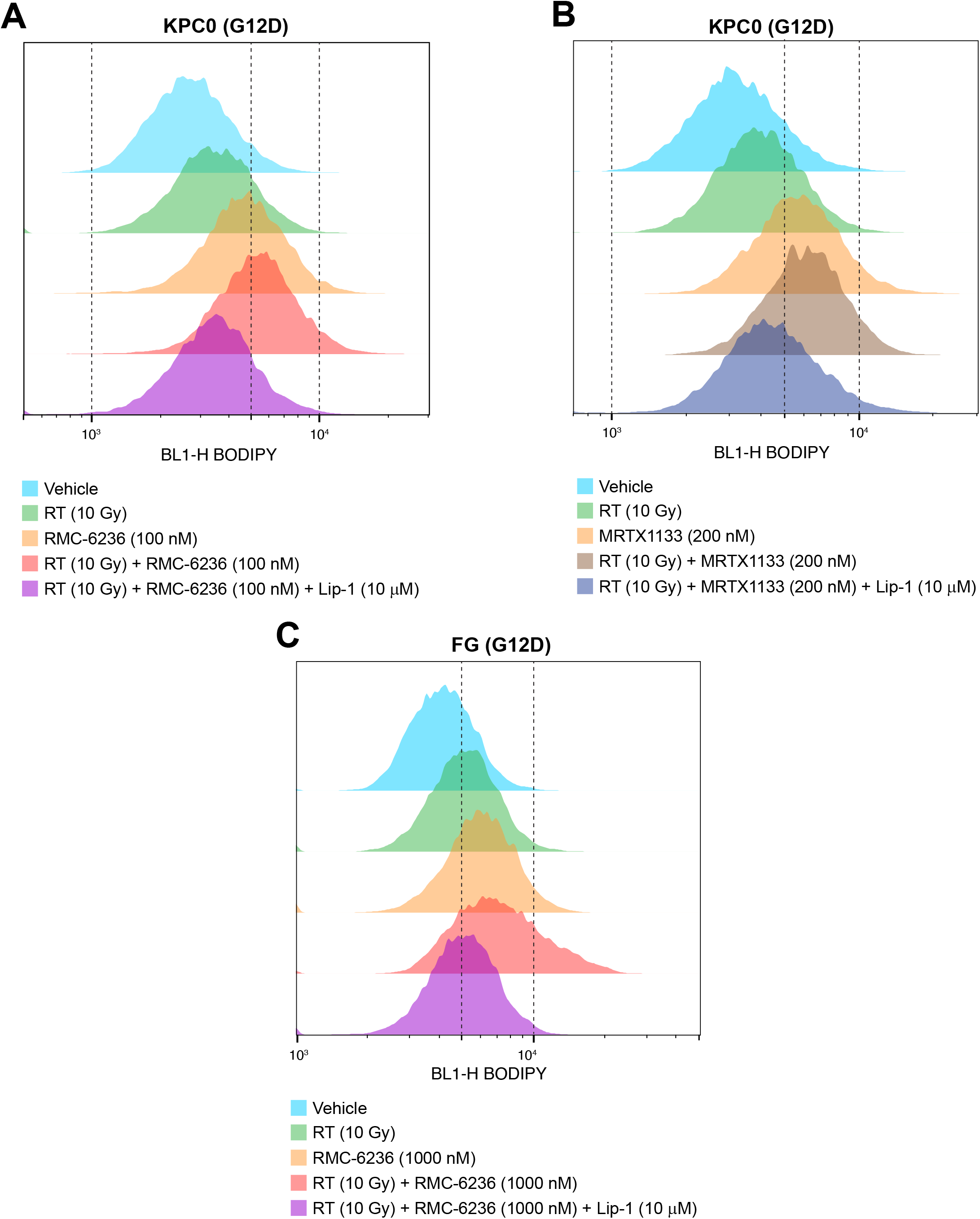
Lipid peroxidation assay. Mouse (KPC0, **A** and **B**) and human (**C**, FG) PDAC cells were treated with vehicle, RT (10 Gy), RMC-6236 (**A** and **C**) or MRTX-1133 (**B**), or the combination then lipid peroxidation was measured by BODIPY C11 581/591 staining and flow cytometry analysis. A ferroptosis/lipid peroxidation inhibitor (liproxstatin-1 or Lip-1) was added to the RT + KRAS inhibitor combination to confirmt the induction of ferroptosis by the treatment.

To evaluate SBRT + KRASi as a therapeutic strategy in a pre-clinical PDAC tumor model, we established syngeneic orthotopic PDAC tumors by injecting luciferase-tagged KPC0 (KPC0-luc) cells into the pancreas of C57BL/6 mice. Similar models have been extensively used to evaluate the therapeutic efficacy of KRASi ^40,42,53-55^. To enable mouse SBRT using the SARRP small animal irradiation platform, we co-transplanted a titanium ligating microclip as a fiducial marker to assist SARRP’s co-registered microCT imaging for localizing the established orthotopic tumors and treatment planning. We utilized an 8 Gy x 5 fraction SBRT treatment schedule to mimic a clinically relevant fractionation schedule for PDAC treatment. Compared to vehicle treated-mice that quickly succumbed to the fast-growing tumor (with a median survival of ∼23 days), SBRT (with a median survival of ∼43 days) or KRASi (with a median survival of ∼65 days) significantly extended the survival. More significantly, the SBRT + KRASi combination treatment achieved long-term tumor control and greatly extended survival (**Fig. 3A and B**). In the SBRT + MRTX1133 group, at the arbitrary endpoint when the longest survival reached 560 days, no tumor relapse had occurred in any of the mice based on post-mortem histopathological examination (**Supplementary Table 1**). Death in the 7 out of 16 mice in this group was likely due to either long-term effects of SBRT or other non-oncologic causes (**Fig. 3A**). However, although SBRT + RMC-6236 similarly extended survival, the majority of mice still had detectable tumors at the time of euthanasia. The few non-tumor related deaths were due to rectal prolapse from unknown causes (**Fig. 3B**). To assess the contribution of enhanced ferroptosis to the improved tumor control and extended survival, we further combined SBRT + KRASi with the ferroptosis inhibitor liproxstatin-1^28,30^. Intriguingly, liproxstatin-1 treatment significantly abolished the enhanced tumor control and survival benefit of SBRT + KRASi. The survival in this group mimicked the survival of the KRASi-only groups (**Fig. 3A and B**). These findings suggest that SBRT + KRASi can synergistically achieve long-term tumor control and greatly extend survival even in the aggressive orthotopic KPC tumor model, and enhanced ferroptosis by this treatment combination contributes to the therapeutic efficacy.

**Figure 3.**
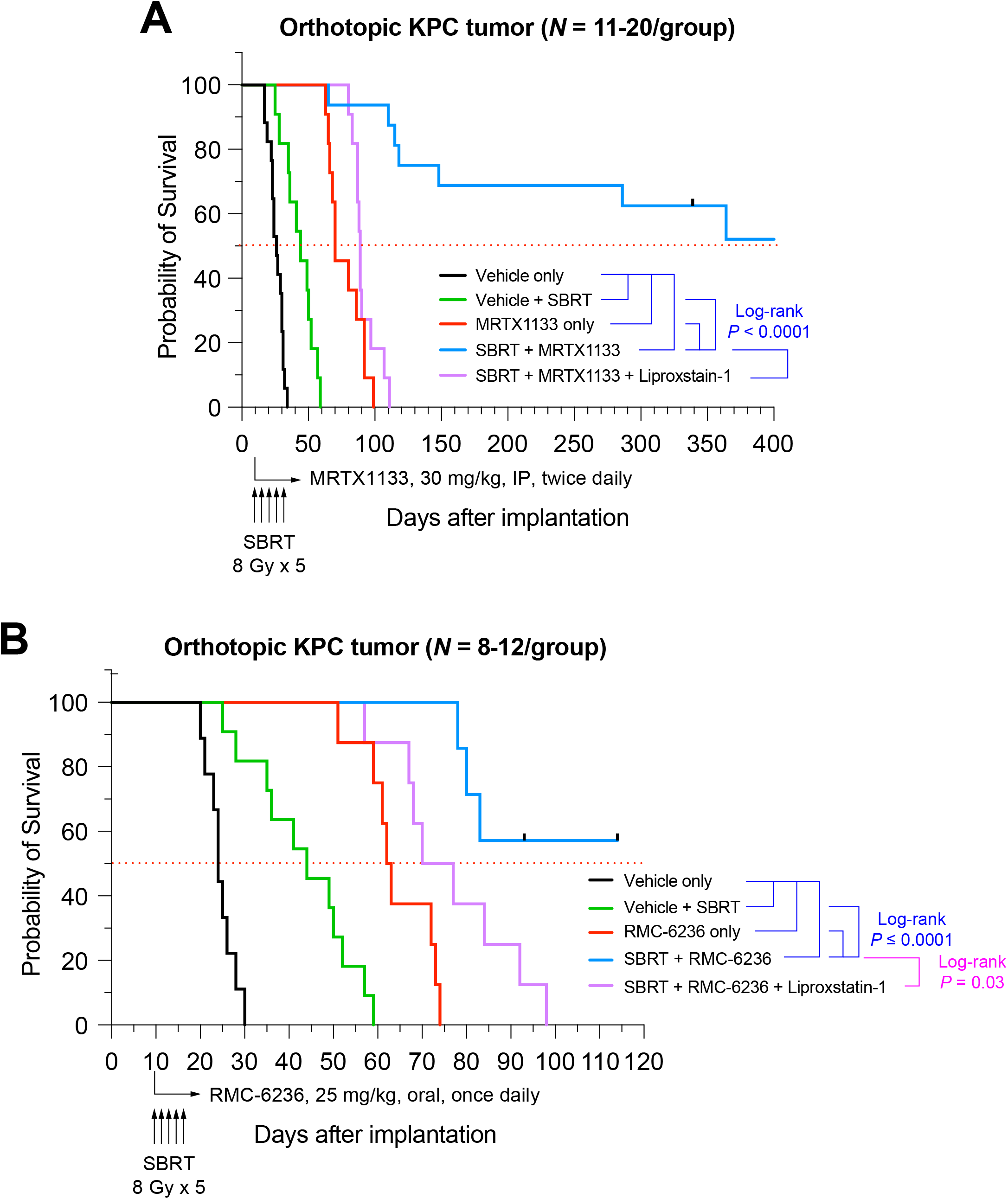
Survival analyses of mice orthotopically transplanted with KPC tumor cells, then treated with SBRT or KRAS inhibitor or the combination. Orthotopic pancreatic tumors were established by injecting luciferase-tagged KPC0 tumor cells (KPC0-luc) into the tail of pancrease in C57BL/6 mice. Once tumor establishment was confirmed by luciferase imaging, the mice were randamized into treatment groups (8-20 per group) to receive vehicle, SBRT (8 Gy x 5), MRTX1133 (**A**) or daraxonrasib (RMC-6236, **B**), or the combination of SBRT and KRAS inhibitor. A ferroptosis inhibitor, liproxstatin-1, was also combined with the combination treatment as a separate group to assess the contribution of ferroptosis to tumor control. The horizontal arrowhead marks the start of vehicle or KRAS inhibitor treatment and the vertial arrowheads show the SBRT fractions. The *P* values of Log-rank tests are also shown.

## DISCUSSION

The development of targeted therapies against mutant KRAS represents a major breakthrough in cancer research, particularly in PDAC, where KRAS mutations are nearly ubiquitous^39^. However, rapid and multifaceted development of resistance to KRAS inhibitors hinders the clinical efficacy of such treatment and limits the improvement in outcome^45^. Resistance to KRAS inhibitors can be broadly classified into primary, acquired, and adaptive mechanisms. (1) Primary resistance refers to the lack of an initial therapeutic response and occurs in approximately 9–24% of patients^56^. (2) Acquired resistance arises following an initial period of tumor regression, typically emerging within months of treatment initiation^46^. One major mechanism of acquired resistance involves on-target alterations in KRAS itself. Secondary mutations within the switch II pocket – the binding site of many KRAS inhibitors – can impair drug binding^57^. In addition, tumors may develop new activating KRAS mutations or amplification of the *KRAS* gene, thereby restoring downstream signaling despite continued inhibition^58^. (3) During adaptive, non-genetic resistance, feedback reactivation of the RTK/MAPK signaling can rapidly restore proliferative signaling^59,60^. Additionally, epithelial–mesenchymal transition (EMT) and other forms of cellular plasticity can give rise to tumor cell states that are less dependent on KRAS signaling^45^. These resistance mechanisms pose significant challenges for PDAC treatment, as KRAS inhibitor monotherapy is unlikely to be sufficient to achieve durable response and combination strategies that target actionable vulnerabilities are required to further improve outcome^45^.

SBRT has emerged as an important modality in the management of PDAC, particularly for patients with locally advanced, borderline resectable, or medically inoperable disease^61^. SBRT delivers highly conformal, ablative doses of radiation in a limited number of fractions, allowing for improved local tumor control while minimizing exposure to surrounding critical structures such as the duodenum and stomach^9,62^. However, definitive chemoradiation to PDAC is challenging because the adjacent stomach and duodenum cannot tolerate high doses of RT^8^. Due to this organ constraint on dose escalation, both intrinsic and acquired resistance of PDAC to RT are commonly observed during the clinical application of SBRT. Therefore, combination strategies that further sensitize PDAC to RT without overt adverse reaction are also needed to improve treatment efficacy and delay or even eliminate tumor relapse.

In the current study, we demonstrated a clear synergism between KRASi and radiation in pre-clinical PDAC models of various origins and mutant *KRAS* alleles. As one of the prominent contributing factors to radiosensitization, KRAS inhibition enhances RT-induced ferroptosis, a form of iron-dependent, non-apoptotic cell death. Since induction of ferroptosis is governed by the various anti-ferroptotic mechanisms, it is likely that KRAS inhibition abrogates or weakens at least some of these cellular defensive mechanisms against ferroptosis. Ferroptosis was initially discovered to be induced by certain compounds in a synthetic lethality fashion in mutant RAS-expressing cells^63-66^. Through activation of ETS-1 in synergy with ATF4, mutant KRAS upregulates SLC7A11 expression and maintains high levels of intracellular cysteine and glutathione (GSH) in cancer cells in response to oxidative stress^67^. Genetic depletion or pharmacological inhibition of SLC7A11 sensitizes mutant KRAS-expressing lung cancer cells to sulfasalazine (a FIN) both in *in vitro* and *in vivo*^67,68^. A recent study showed that through downstream activation of MAPK in synergy with NRF2, mutant KRAS also upregulates the expression of ferroptosis suppressor protein 1 (FSP1) to mediate ferroptosis resistance and pharmacological FSP1 inhibition sensitized KRAS^G12D^-driven mouse PDAC organoids to RAS-selective lethal 3 (RSL3)-induced ferroptosis^69^. Mutant KRAS also upregulates the expression of lipid metabolism genes such as acyl-coenzyme A synthetase long chain family member 3 (*ACSL*3) and fatty acid synthase (*FASN*), which promotes saturated fatty acid (SFA) and monounsaturated fatty acid (MUFA) biosynthesis and ferroptosis resistance in lung cancer cells^70-72^. The specific KRAS-mediated anti-ferroptotic mechanisms that contribute to radioresistance in PDAC cells are pending further investigation.

In conclusion, resistance to KRAS inhibitors and RT in pancreatic cancer is a multifaceted and dynamic process driven by genetic alterations, signaling rewiring, and cellular plasticity. The complexity and heterogeneity of these resistance mechanisms necessitate combination approaches that target multiple pathways simultaneously. Advances in the development of next-generation KRAS inhibitors, together with improved understanding of resistance biology, provide a strong foundation for the design of more effective therapeutic strategies. Ultimately, the successful integration of targeted therapy, combination regimens, and biomarker-driven patient selection will be critical for achieving meaningful and durable clinical benefit in patients with KRAS-mutant PDAC.

## ACKNOWLEDGEMENTS

This research was supported by the Olga Keith and Harry Carothers Wiess Distinguished University Chair in Cancer Research Endowment Fund (to A.C.K.), National Institutes of Health grants U54CA274220 (to A.C.K., D.J. and B.G.), R01CA269646 (to B.G.), and R01CA301148 (to B.G.), the Collaborative Accelerator for Transformative Research Endeavors grant, jointly awarded by The University of Texas at Austin and The University of Texas MD Anderson Cancer Center (to B.G.), and the N.G. and Helen T. Hawkins Distinguished Professorship for Cancer Research of MD Anderson Cancer Center (to B.G.).

## AUTHOR CONTRIBUTIONS

D.J. and A.C.K. designed and supervised the study. T.W. performed the majority of the *in vitro* and *in vivo* experiments, with technical assistance from L.W., J.X., Y.G., and L.X. D.J. and T.W. analyzed the data and wrote the manuscript. Y.L. and F.G. contributed to the design of the *in vitro* radiation treatment and radiation dosimetry measurements for both *in vitro* and *in vivo* studies. B.G., D.H. and V.B. contributed substantially to discussion of the content and reviewed and/or edited the manuscript before submission.

## REFERENCES

1. Siegel, R.L., Giaquinto, A.N. & Jemal, A. Cancer statistics, 2024. CA Cancer J Clin 74, 12–49 (2024).

2. Rahib, L., Wehner, M.R., Matrisian, L.M. & Nead, K.T. Estimated Projection of US Cancer Incidence and Death to 2040. JAMA Netw Open 4, e214708 (2021).

3. Ryan, D.P., Hong, T.S. & Bardeesy, N. Pancreatic adenocarcinoma. The New England journal of medicine 371, 1039–1049 (2014).

4. Bengtsson, A., Andersson, R. & Ansari, D. The actual 5-year survivors of pancreatic ductal adenocarcinoma based on real-world data. Sci Rep 10, 16425 (2020).

5. Li, D., Xie, K., Wolff, R. & Abbruzzese, J.L. Pancreatic cancer. Lancet 363, 1049–1057 (2004).

6. Hattangadi, J.A., Hong, T.S., Yeap, B.Y. & Mamon, H.J. Results and patterns of failure in patients treated with adjuvant combined chemoradiation therapy for resected pancreatic adenocarcinoma. Cancer 115, 3640–3650 (2009).

7. Balaban, E.P., Mangu, P.B. & Yee, N.S. Locally Advanced Unresectable Pancreatic Cancer: American Society of Clinical Oncology Clinical Practice Guideline Summary. J Oncol Pract 13, 265–269 (2017).

8. Kelly, P., et al. Duodenal toxicity after fractionated chemoradiation for unresectable pancreatic cancer. Int J Radiat Oncol Biol Phys 85, e143–149 (2013).

9. Koong, A.C., et al. Phase I study of stereotactic radiosurgery in patients with locally advanced pancreatic cancer. Int J Radiat Oncol Biol Phys 58, 1017–1021 (2004).

10. Palta, M., et al. Radiation Therapy for Pancreatic Cancer: Executive Summary of an ASTRO Clinical Practice Guideline. Pract Radiat Oncol 9, 322–332 (2019).

11. Delpu, Y., et al. Genetic and epigenetic alterations in pancreatic carcinogenesis. Curr Genomics 12, 15–24 (2011).

12. Bournet, B., Buscail, C., Muscari, F., Cordelier, P. & Buscail, L. Targeting KRAS for diagnosis, prognosis, and treatment of pancreatic cancer: Hopes and realities. Eur J Cancer 54, 75–83 (2016).

13. Haigis, K.M. KRAS Alleles: The Devil Is in the Detail. Trends Cancer 3, 686–697 (2017).

14. di Magliano, M.P. & Logsdon, C.D. Roles for KRAS in pancreatic tumor development and progression. Gastroenterology 144, 1220–1229 (2013).

15. Jonckheere, N., Vasseur, R. & Van Seuningen, I. The cornerstone K-RAS mutation in pancreatic adenocarcinoma: From cell signaling network, target genes, biological processes to therapeutic targeting. Crit Rev Oncol Hematol 111, 7–19 (2017).

16. Dixon, S.J., et al. Ferroptosis: an iron-dependent form of nonapoptotic cell death. Cell 149, 1060–1072 (2012).

17. Stockwell, B.R., et al. Ferroptosis: A Regulated Cell Death Nexus Linking Metabolism, Redox Biology, and Disease. Cell 171, 273–285 (2017).

18. Mao, C., Jiang, D., Koong, A.C. & Gan, B. Exploiting metabolic cell death for cancer therapy. Nat Rev Cancer 26, 27–45 (2026).

19. Conrad, M. & Pratt, D.A. The chemical basis of ferroptosis. Nat Chem Biol 15, 1137–1147 (2019).

20. Seibt, T.M., Proneth, B. & Conrad, M. Role of GPX4 in ferroptosis and its pharmacological implication. Free Radic Biol Med 133, 144–152 (2019).

21. Friedmann Angeli, J.P., et al. Inactivation of the ferroptosis regulator Gpx4 triggers acute renal failure in mice. Nature cell biology 16, 1180–1191 (2014).

22. Yang, W.S., et al. Regulation of ferroptotic cancer cell death by GPX4. Cell 156, 317–331 (2014).

23. Bersuker, K., et al. The CoQ oxidoreductase FSP1 acts parallel to GPX4 to inhibit ferroptosis. Nature 575, 688–692 (2019).

24. Doll, S., et al. FSP1 is a glutathione-independent ferroptosis suppressor. Nature 575, 693–698 (2019).

25. Mao, C., et al. DHODH-mediated ferroptosis defence is a targetable vulnerability in cancer. Nature 593, 586–590 (2021).

26. Lei, G., Zhuang, L. & Gan, B. Targeting ferroptosis as a vulnerability in cancer. Nat Rev Cancer 22, 381–396 (2022).

27. Lang, X., et al. Radiotherapy and Immunotherapy Promote Tumoral Lipid Oxidation and Ferroptosis via Synergistic Repression of SLC7A11. Cancer Discov 9, 1673–1685 (2019).

28. Lei, G., et al. The role of ferroptosis in ionizing radiation-induced cell death and tumor suppression. Cell Res 30, 146–162 (2020).

29. Ye, L.F., et al. Radiation-Induced Lipid Peroxidation Triggers Ferroptosis and Synergizes with Ferroptosis Inducers. ACS Chem Biol (2020).

30. Lei, G., et al. Ferroptosis as a mechanism to mediate p53 function in tumor radiosensitivity. Oncogene 40, 3533–3547 (2021).

31. Chen, X., Kang, R., Kroemer, G. & Tang, D. Targeting ferroptosis in pancreatic cancer: a double-edged sword. Trends Cancer 7, 891–901 (2021).

32. Badgley, M.A., et al. Cysteine depletion induces pancreatic tumor ferroptosis in mice. Science 368, 85–89 (2020).

33. Kremer, D.M., et al. GOT1 inhibition promotes pancreatic cancer cell death by ferroptosis. Nat Commun 12, 4860 (2021).

34. Cox, A.D., Fesik, S.W., Kimmelman, A.C., Luo, J. & Der, C.J. Drugging the undruggable RAS: Mission possible? Nat Rev Drug Discov 13, 828–851 (2014).

35. Fell, J.B., et al. Identification of the Clinical Development Candidate MRTX849, a Covalent KRAS(G12C) Inhibitor for the Treatment of Cancer. J Med Chem 63, 6679–6693 (2020).

36. Lanman, B.A., et al. Discovery of a Covalent Inhibitor of KRAS(G12C) (AMG 510) for the Treatment of Solid Tumors. J Med Chem 63, 52–65 (2020).

37. Singhal, A., Li, B.T. & O’Reilly, E.M. Targeting KRAS in cancer. Nat Med 30, 969–983 (2024).

38. Wang, X., et al. Identification of MRTX1133, a Noncovalent, Potent, and Selective KRAS(G12D) Inhibitor. J Med Chem 65, 3123–3133 (2022).

39. Riedl, J.M., et al. Emerging landscape of KRAS inhibitors in cancer treatment. Cancer Cell 44, 471–497 (2026).

40. Kemp, S.B., et al. Efficacy of a Small-Molecule Inhibitor of KrasG12D in Immunocompetent Models of Pancreatic Cancer. Cancer Discov 13, 298–311 (2023).

41. Hallin, J., et al. Anti-tumor efficacy of a potent and selective non-covalent KRAS(G12D) inhibitor. Nat Med 28, 2171–2182 (2022).

42. Mahadevan, K.K., et al. KRAS(G12D) inhibition reprograms the microenvironment of early and advanced pancreatic cancer to promote FAS-mediated killing by CD8(+) T cells. Cancer Cell 41, 1606–1620 e1608 (2023).

43. Mahadevan, K.K., et al. Elimination of oncogenic KRAS in genetic mouse models eradicates pancreatic cancer by inducing FAS-dependent apoptosis by CD8(+) T cells. Dev Cell 58, 1562–1577 e1568 (2023).

44. O’Reilly, E.M., et al. Daraxonrasib or Chemotherapy in Previously Treated Metastatic Pancreatic Cancer. The New England journal of medicine (2026).

45. Isermann, T., Sers, C., Der, C.J. & Papke, B. KRAS inhibitors: resistance drivers and combinatorial strategies. Trends Cancer 11, 91–116 (2025).

46. Awad, M.M., et al. Acquired Resistance to KRAS(G12C) Inhibition in Cancer. The New England journal of medicine 384, 2382–2393 (2021).

47. Dilly, J., et al. Mechanisms of Resistance to Oncogenic KRAS Inhibition in Pancreatic Cancer. Cancer Discov 14, 2135–2161 (2024).

48. Vezeridis, M.P., et al. In vivo selection of a highly metastatic cell line from a human pancreatic carcinoma in the nude mouse. Cancer 69, 2060–2063 (1992).

49. Kim, M.P., et al. Generation of orthotopic and heterotopic human pancreatic cancer xenografts in immunodeficient mice. Nat Protoc 4, 1670–1680 (2009).

50. Jiang, D., et al. IRE1alpha determines ferroptosis sensitivity through regulation of glutathione synthesis. Nat Commun 15, 4114 (2024).

51. Zheng, S., et al. SynergyFinder Plus: Toward Better Interpretation and Annotation of Drug Combination Screening Datasets. Genomics Proteomics Bioinformatics 20, 587–596 (2022).

52. Berenbaum, M.C. What is synergy? Pharmacol Rev 41, 93–141 (1989).

53. McAndrews, K.M., et al. An allele-agnostic mutant-KRAS inhibitor suppresses tumor maintenance signals and reprograms tumor immunity in pancreatic cancer. Sci Transl Med 17, eadt5511 (2025).

54. Mahadevan, K.K., et al. Distinct small molecule inhibitors of Kras specifically prime CTLA4 blockade therapy to transcriptionally reprogram Tregs and overcome resistance to suppress pancreas cancer. bioRxiv (2025).

55. Wasko, U.N., et al. Tumour-selective activity of RAS-GTP inhibition in pancreatic cancer. Nature 629, 927–936 (2024).

56. Hong, D.S., et al. KRAS(G12C) Inhibition with Sotorasib in Advanced Solid Tumors. The New England journal of medicine 383, 1207–1217 (2020).

57. Tanaka, N., et al. Clinical Acquired Resistance to KRAS(G12C) Inhibition through a Novel KRAS Switch-II Pocket Mutation and Polyclonal Alterations Converging on RAS-MAPK Reactivation. Cancer Discov 11, 1913–1922 (2021).

58. Zhao, Y., et al. Diverse alterations associated with resistance to KRAS(G12C) inhibition. Nature 599, 679–683 (2021).

59. Prahallad, A., et al. Unresponsiveness of colon cancer to BRAF(V600E) inhibition through feedback activation of EGFR. Nature 483, 100–103 (2012).

60. Corcoran, R.B., et al. EGFR-mediated re-activation of MAPK signaling contributes to insensitivity of BRAF mutant colorectal cancers to RAF inhibition with vemurafenib. Cancer Discov 2, 227–235 (2012).

61. Burkon, P., et al. Stereotactic Body Radiotherapy (SBRT) of Pancreatic Cancer-A Critical Review and Practical Consideration. Biomedicines 10(2022).

62. Herman, J.M., et al. Phase 2 multi-institutional trial evaluating gemcitabine and stereotactic body radiotherapy for patients with locally advanced unresectable pancreatic adenocarcinoma. Cancer 121, 1128–1137 (2015).

63. Dixon, S.J. & Stockwell, B.R. The Hallmarks of Ferroptosis. Annual Review of Cancer Biology 3, 35–54 (2019).

64. Dolma, S., Lessnick, S.L., Hahn, W.C. & Stockwell, B.R. Identification of genotype-selective antitumor agents using synthetic lethal chemical screening in engineered human tumor cells. Cancer Cell 3, 285–296 (2003).

65. Yagoda, N., et al. RAS-RAF-MEK-dependent oxidative cell death involving voltage-dependent anion channels. Nature 447, 864–868 (2007).

66. Yang, W.S. & Stockwell, B.R. Synthetic lethal screening identifies compounds activating iron-dependent, nonapoptotic cell death in oncogenic-RAS-harboring cancer cells. Chem Biol 15, 234–245 (2008).

67. Lim, J.K.M., et al. Cystine/glutamate antiporter xCT (SLC7A11) facilitates oncogenic RAS transformation by preserving intracellular redox balance. Proc Natl Acad Sci U S A 116, 9433–9442 (2019).

68. Hu, K., et al. Suppression of the SLC7A11/glutathione axis causes synthetic lethality in KRAS-mutant lung adenocarcinoma. J Clin Invest 130, 1752–1766 (2020).

69. Muller, F., et al. Elevated FSP1 protects KRAS-mutated cells from ferroptosis during tumor initiation. Cell Death Differ 30, 442–456 (2023).

70. Magtanong, L., et al. Exogenous Monounsaturated Fatty Acids Promote a Ferroptosis-Resistant Cell State. Cell Chem Biol 26, 420–432 e429 (2019).

71. Padanad, M.S., et al. Fatty Acid Oxidation Mediated by Acyl-CoA Synthetase Long Chain 3 Is Required for Mutant KRAS Lung Tumorigenesis. Cell Rep 16, 1614–1628 (2016).

72. Bartolacci, C., et al. Targeting de novo lipogenesis and the Lands cycle induces ferroptosis in KRAS-mutant lung cancer. Nat Commun 13, 4327 (2022).

